# A Reinforcement Meta-Learning Framework of Executive Function and Information Demand

**DOI:** 10.1101/2021.07.18.452793

**Authors:** Massimo Silvetti, Stefano Lasaponara, Mattias Horan, Jacqueline Gottlieb

## Abstract

Gathering information is crucial for maximizing fitness, but requires diverting resources from searching directly for primary rewards to actively exploring the environment. Optimal decision-making thus maximizes information while reducing effort costs, but little is known about the neural implementation of these tradeoffs. We present a Reinforcement Meta-Learning (RML) computational mechanism that solves the trade-offs between the value and costs of gathering information. We implement the RML in a biologically plausible architecture that links catecholaminergic neuromodulators, the medial prefrontal cortex and topographically organized visual maps and show that it accounts for neural and behavioral findings on information demand motivated by instrumental incentives and intrinsic utility. Moreover, the utility function used by the RML, encoded by dopamine, is an approximation of free-energy. Thus, the RML presents a biologically plausible mechanism through which coordinated motivational, executive and sensory systems generate visual information gathering policies that minimize free energy.

## Introduction

Reducing uncertainty about the state of the world is a fundamental imperative for biological organisms. To efficiently meet this imperative, animals must balance the benefits of taking actions that obtain information (reduce uncertainty) against the costs that these actions entail. Converging evidence suggests that core processes serving information gathering are attention and active sensing behaviors that sample rich sensory streams. Computational and behavioral studies in humans and monkeys show that visual active sensing using rapid eye movements (saccades) partly conform with optimal information gathering including the minimization of “free-energy”^1-3^. However, we lack a neurocomputational account of how this process emerges in a biologically plausible attention control architecture.

Theories of executive function propose that the allocation of cognitive resources relies on the interaction between a “monitoring” process implemented in the medial prefrontal cortex (MPFC), and “regulatory” mechanisms implemented in posterior and lateral areas^4-6^. The MPFC circuit estimates the utility of a task and the effort to invest in the task, while regulatory networks implement the selected cognitive policy.

We recently proposed a computational model of the monitoring role of the MPFC using a reinforcement metal learner (RML) mechanism^7^. The RML is a bio-inspired autonomous agent that maximizes long-term rewards while minimizing costs^8^. In the service of reward maximization, the RML optimizes not only overt motor behavior but also *internal* neural dynamics that indirectly influence behavior. The optimization of internal dynamics is referred to as “meta-learning” and modulates cognitive functions in pursuit of a goal^9,10^. In our RML model, the MPFC receives reward information through dopaminergic (DA) afferents from the ventral tegmental area (VTA) and calls for a *boost* of norepinephrine from the LC. The NE boost controls internal dynamics to enhance relevant information processing, but it registers as a *cost* and is thus limited by the extent to which it enhances utility. In a recent report we showed that this general-purpose optimization mechanism accounts for neural and behavioral findings in independent domains including adjustments of learning rates, physical effort, working memory, and higher order conditioning^7^.

Here we show that the same architecture accounts for neural and behavioral findings on attentional information gathering. We simulate *instrumental* information demand, in which agents gather visual information for guiding an incentivized choice, and *non-instrumental* information demand, in which they seek information for its own sake. We show that, when connected with a topographic visual map, the RML reproduces the distinct responses to expected information gains and expected reward gains that have been described in the lateral intraparietal area (LIP)^11^ and support a new interpretation of attentional priority as a cognitive state that is optimized for reducing uncertainty. We next show that, without an increase in model complexity, the RML reproduces distinct informational drives related to the intrinsic desire to reduce uncertainty versus the desire to anticipate positive outcomes^2^. Importantly, we show that these operations implement the free-energy principle^12^ and are thus a biologically plausible theory on how the brain generates uncertainty reduction policies in response to external incentives and intrinsic utility.

## Results

### The Reinforcement Meta-Learner (RML)

The RML^7^ is a bio-inspired autonomous agent that optimizes long-term rewards using four interacting modules (**Fig. 1A**). Two modules are subcortical and represent the VTA and LC (**Fig. 1A**, red and orange). The VTA module simulates the release of DA that signals extrinsic and intrinsic *rewards* (see below). The LC module simulates release of NE and signals cognitive or physical effort^13,14^ (see *Discussion* for further justification of this choice). Two additional cortical modules - the MPFC_Act_ and MPFC_Boost_ – represent the MPFC (**Fig. 1A**, blue and green). These modules use reward and effort information from the VTA and LC along with state information from the external environment to update state-action values and select appropriate actions. The MPFC_Act_ module selects motor actions – i.e., makes decisions that act on the external space. The MPFC_Boost_ module makes decisions oriented toward the internal space by generating control signals that call for *boosts* of NE^15-17^. Boosting NE promotes effortful actions (e.g., by modulating the motor decisions made by the MPFC_act_) and enhances information processing in other brain structures, as we explain next. The boost, however, is registered as a cost, and the MPFC_boost_ module implements meta-learning by dynamically optimizing the trade-off between this *cost* and the *rewards* associated with boosting (see Methods, eq. 5, 6, 8b).

**Figure 1.**
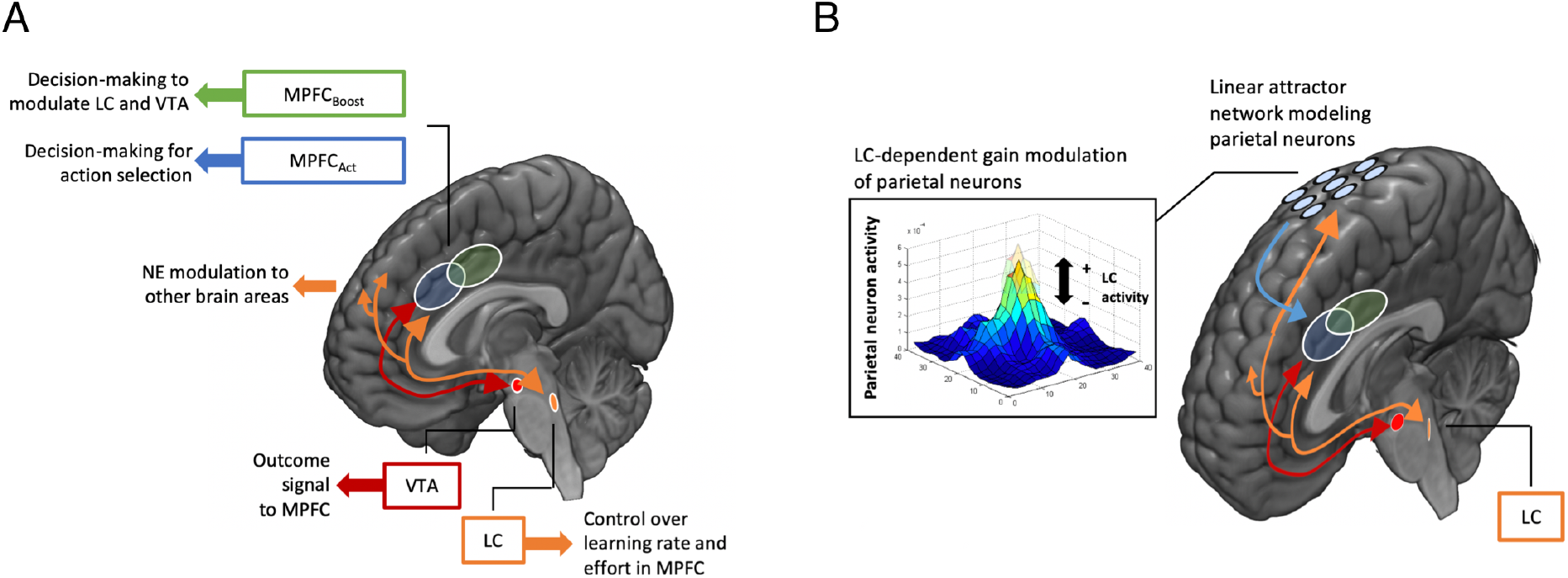
Overview of the RML with mapping on brain structures. **A**) The RML consists of two modules conveying information about rewards from the ventral tegmental area (VTA) and effort from the locus ceruleus (LC) that target the two modules of the medial prefrontal cortex (MPFC). The MFPC modules perform action-outcome comparison to optimize state-action selection involving motor actions that affect the external environment (MPFC_Act_) and internal actions that modulate the activity of VTA and LC (MPFC_Boost_). **B**) The hybrid attention-RML (aRML) connects the core RML in **A** to a map of visual space simulating topographically organized visual areas. The visual map is implemented as a linear attractor basis function network integrating retinal position and current eye position^18^ and is connected to the RML through an efferent copy of the LC output that enhances visual response gains (orange arrow). Information about saccade target selection is directed toward the MPFC_Act_ module, improving decisionmaking to obtain rewards (blue arrow). Panel **A** is modified with permission from ^7^.

We previously showed that the RML is a general-purpose optimizer that can use LC-dependent boost signals to regulate the activity of computational modules implementing memory, learning rates, or cognitive effort^7,19-21^. In the following sections, we describe how the RML acts when connected with systems of saccade and attention in tasks of information demand.

### Optimizing visual priority for instrumental information

The selection of targets for saccades and attention is mediated by a network of retinotopically organized structures that include mid-level visual areas, fronto-parietal areas and the superior colliculus^22^. We recently showed that activity in one of these areas – area LIP - depends on expected information gains during a task of instrumental information demand^11^. To understand the neuro-computational mechanisms of these findings, we connected the RML to a neural network for visual space representation, creating a hybrid model we refer to as the *attention-RML* (aRML, **Fig. 1B**).

Consistent with neurophysiological studies, the visual network has spatially tuned visual receptive fields (RF) and maintains spatial constancy across shifts of gaze (modeled through basis function coordinate transformation^18^. In the aRML, the visual map receives LC/NE output from the MPFC-VTA-LC circuit, which enhances sensory gains and improves performance for saccades and attention^23,24^ (**Fig. 1B**, orange arrow). Topographic visual information is then redirected to the MPFC_Act_ and used to make a decision (**Fig. 1B**, blue arrow; see *Methods* for a detailed description of the aRML).

To investigate how the visual map responds during information demand, we administered the same task to the aRML that we used in the non-human primates. The task required the agent (monkey or aRML) to make an instrumental decision, choosing one of two targets to obtain a reward (**Fig. 2A**, “Final Decision”). Before making this choice, the agent directed gaze to a cue (**Fig. 2A**, “Cue”) which triggered relevant information consisting of visual motion toward the correct decision alternative (**Fig. 2A**, “Final Decision”). We examined how the visual responses to the informative cue changed in response to the information gains and reward gains the cue was expected to bring, which we orthogonally manipulated (**Fig. 2A**, right). We manipulated information gains using two contexts (trial blocks) that differed in their *ex ante* decision uncertainty. In the “Informative” context, the two decision alternatives had equal prior probability of being correct and the agent could expect that the motion would resolve this uncertainty by indicating the correct alternative on that trial. In the “Uninformative” context, the correct decision alternative was fixed across trials; thus, the agent had no prior uncertainty and could expect that the motion information, while valid, would be redundant with their prior expectation. We manipulated reward gains by randomly interleaving, in both Informative and Uninformative blocks, trials that delivered a small or large reward for a correct choice, signaling reward size at the start of each trial. This created a 2 x 2 task design that orthogonally manipulated the reward and information gains that the agent could expect to experience after making a saccade to obtain the Cue’s information (**Fig. 2A**, right).

**Figure 2.**
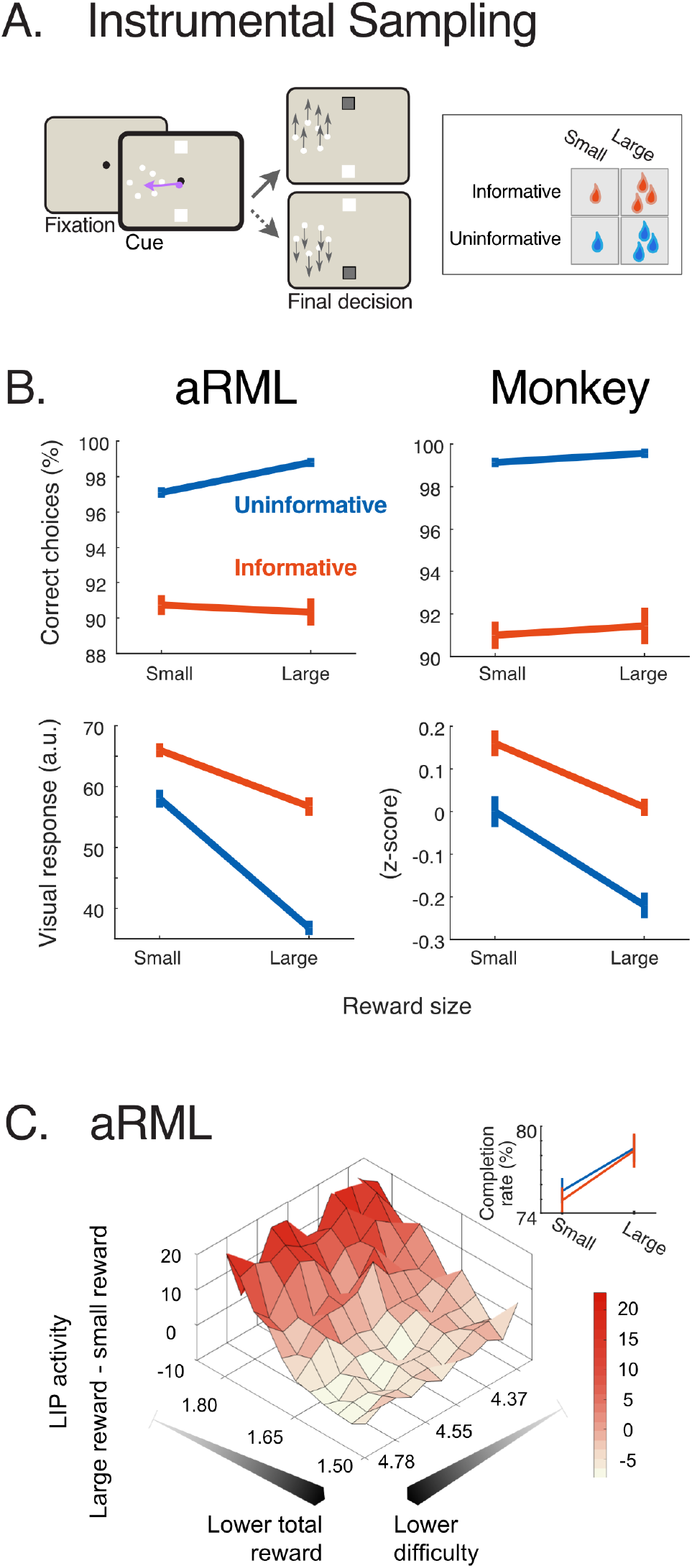
Instrumental Information Sampling Task. **A)** The agent (aRML or monkey) was exposed to a display with two targets (white squares) and an information source (cloud of dots; “Cue”). To obtain a reward, the aRML had to generate a saccade to the cue, trigger motion information and, based on the information, make a correct final choice (choose the target that was congruent with the motion direction). This design was presented in 4 conditions that crossed 2 levels of information gains and 2 reward magnitudes. In informative trial blocks, either target had equal prior probability of being correct and the motion resolved the uncertainty. In uninformative blocks, a single trial was correct on all trials, making the motion information redundant. A correct response in either block could deliver a small or large reward (signaled at the start of each trial). No reward was delivered if the aRML chose the wrong target or chose not to engage in a trial (“break fixation”). **B)** aRML-monkey comparison. Percentage of correct choices (top row) and visual activity (lower row), as a function of reward magnitude in the informative (red) and uninformative (blue) blocks. Points show mean ± s.e.m. over model simulations (left column) or empirical data from^11^ (right column). In the bottom row, the left plot shows visual activation in the aRML in arbitrary units and the right plot shows LIP activity (z-scored firing rates) from^11^. **C)** Reward modulations predicted by the aRML. The y axis shows the difference in activity between large reward and small reward trials as a function of the total reward size and task difficulty (precision of LIP information coding, measured as Fisher Information (I)). For small average rewards and low difficulty, activity is higher for small relative to large rewards, as found in^11^. Inset) The aRML has a lower completion rate (higher probability of aborting a trial) for when smaller rewards are at stake for both informative (red) and uninformative (blue) blocks as found for monkeys in^11^.

We implemented the RML using the standard parameter values^7^ without fitting to empirical observations. For each condition we simulated a number of iterations comparable to the size of empirical data sets, avoiding artificially inflating p-values by increasing the number of iterations^7^ (see *Methods*).

The aRML simulations replicated empirical findings (**Fig. 2B**). The aRML produced lower decision accuracy on informative versus uninformative trials, showing that gathering information is more challenging than acting merely on one’s priors (**Fig. 2B**, top left; F(1,29) = 212.9, p < 0.0001), consistent with behavioral findings (**Fig. 2B**, top right; cf^11^). Critically, the aRML simulated the uncertainty-related enhancement of the visual map, with higher visual responses in informative relative to uninformative trials (**Figure 2B**, bottom left; main effect of informativeness, F(1,29)=89.91, p < 0.0001, n = 30 simulations), replicating the uncertainty-related enhancement in LIP cells (**Fig. 2B**, bottom right).

It is important to note that the aRML generated uncertainty-related enhancement without including uncertainty reduction in the utility function. In its attempt to maximize reward gains, the model finds that a boost of NE enhances reward rates by enhancing attentional priority, which in turn improves discrimination accuracy and, ultimately, decision accuracy. Because the model registers the NE boost as a *cost*, it deploys it selectively, only in informative trials when the information enhances reward expectations. Thus, in instrumental conditions, uncertainty-related enhancement can emerge strictly as a consequence of reward maximization; however, in the following section, we show that this process can be enhanced by an explicit drive for reducing uncertainty.

A striking empirical finding was that LIP cells responded to reward magnitudes with a *negative* modulation – showing stronger responses when a smaller rather than larger reward was at stake (**Fig. 2B**, bottom right). The aRML replicated this finding (**Fig. 2B**, bottom left) and suggested that it was a compensatory mechanism triggered at specific combinations of reward size and difficulty. This was shown by simulating the task at a wider range of reward magnitudes and visual discrimination difficulty (**Fig. 2C**). For most of the range shown in **Fig. 2C**, the aRML produced a higher boost of NE and, consequently, higher visual activity when a larger reward was at stake consistent with the typically observed positive reward modulations of the visual map (**Fig. 2C**, red). However, this trend reversed for a set of conditions similar to those used by Horan et al^11^, which had low discrimination difficulty (high motion coherence) and low trial-by-trial rewards (**Fig. 2C**, bottom left corner, white showing negative values). This was due to the fact that the aRML could choose to disengage from the task and did so more frequently when the rewards were low, similar to behavioral findings in monkeys^11^ (**Fig. 2C**, inset; rate of trial completion increases with expected reward; F(1,29) = 14.1, p < 0.0001). To counteract the loss of reward due to incomplete trials, the aRML generated an additional boost of NE. This prevented the rate of trial completion from becoming too low but produced the counterintuitive enhancement at small relative to large reward sizes.

In sum, the aRML reproduces both the uncertainty and reward modulations found in visual maps during instrumental information demand as adaptations of internal dynamics that optimize reward rates given the constraints of a task.

### Intrinsically motivated information seeking in non-instrumental tasks

The aRML described in the previous section generates information demand as a consequence of maximizing instrumental rewards, but humans and monkeys also seek information as a good in itself, independently of external incentives^25^. In this section we show that, without introducing new parameters, the RML reproduces distinct components of non-instrumental information demand related to uncertainty reduction versus gathering positive observations.

The original RML estimated uncertainty as expected surprisal (to produce volatility-based adjustment of learning rates (^7^; and Methods, eq. 7d), and here we simply included this term in the function defining DA activity. The utility function that is conveyed by DA in this modified model (which we call the RML-C, for RML-Curiosity; eq. 11) is written below in a simplified form:

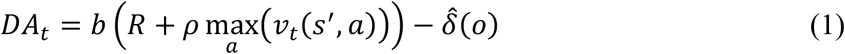

The left-hand side term in this equation is the one used by the original RML and is governed by the current extrinsic reward, R, combined with the value of expected future states, *v (s’,a)* weighted by *ρ*, the temporal discount factor. The right-hand side term, 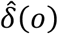, is the expected surprisal - unsigned prediction error of the outcome over trials - a measure of uncertainty correlated with entropy^26,27^. As we explain in the *Discussion*, introducing this term in the loss function represents an implementation of the free-energy principle^12^ in the framework of meta-RL.

We next used both the original RML and the RML-C to simulate behavior in tasks of noninstrumental information demand. We simulated the conditions used in ^2^ in which monkeys could seek information under different prior probabilities of delivering rewards (**Fig. 3A**). Uncovering information was effortful, as before, but it had no instrumental incentives, as the rewards were delivered non-contingently - i.e., were identical whether or not the agent obtained the information (**Fig. 3Ai**).

**Figure 3.**
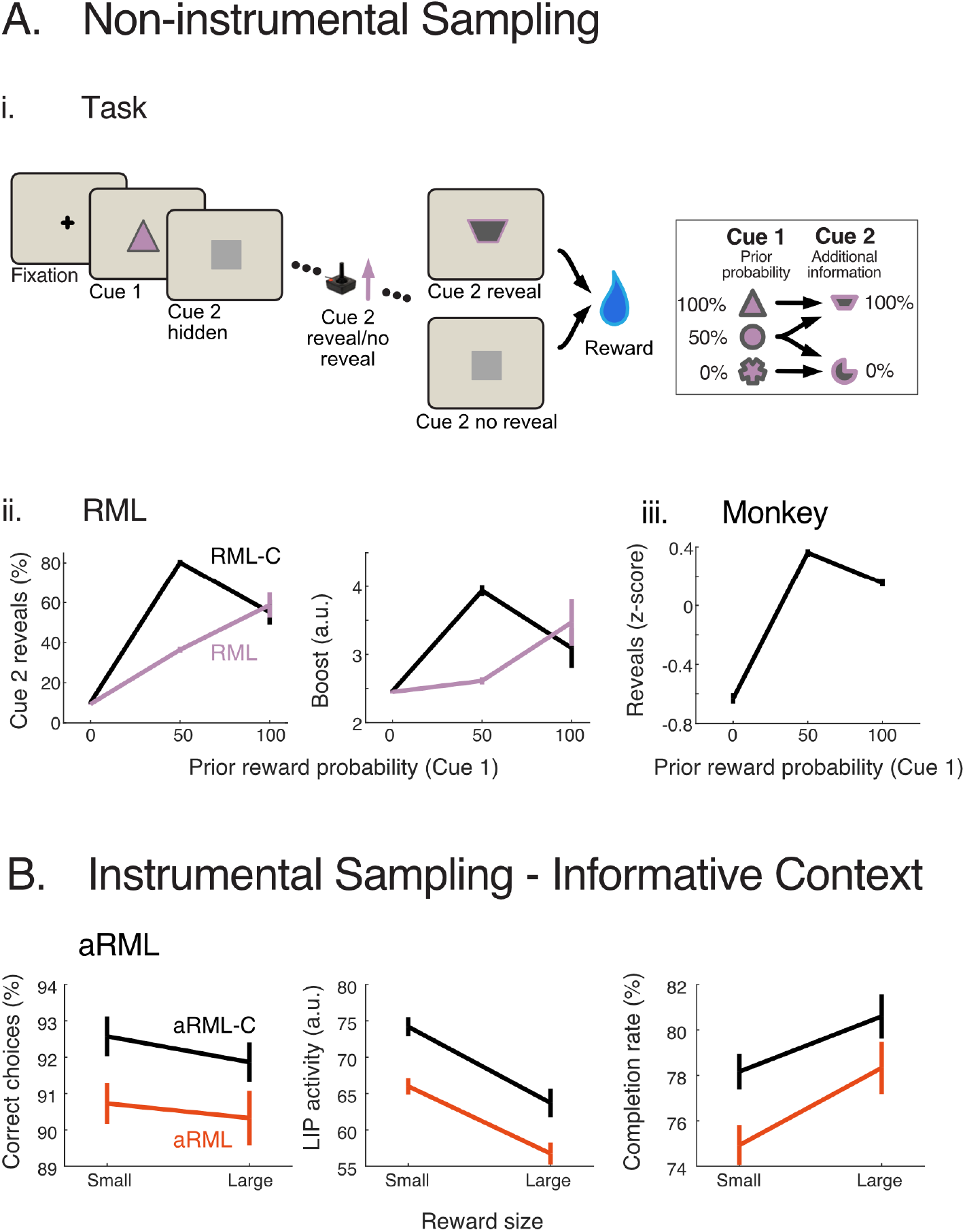
**A. Non-Instrumental Sampling Task. i)** Trial structure. After viewing the prior reward probability (Cue 1), the RML receives a visual mask and can choose to exert effort to remove the mask and uncover Cue 2 or just wait for reward delivery. After a fixed time delay, a reward is delivered based on the cue-reward associations, regardless the decision about uncovering Cue 2. Cue 1 could signal 0%, 50% or 100% prior reward probability. Cue 2 provided perfect information about presence or absence of reward, which was redundant if it followed a 0% and 100% Cue 1 and resolved uncertainty after a 50% Cue 1. **Aii)** Probability of revealing cue 2 (left) and NE boost (right) from the aRML and RML-C as a function of Cue 1 probability. Each point shows mean± s.e.m. over a set of iterations. **Aiii)** Probability of revealing Cue 2 shown by monkeys in^2^. **B) Non-Instrumental Sampling Task.** Predictions of the aRML and aRML-C regarding %correct, LIP activity and completion rate for the informative condition in the Instrumental Information Sampling Task (**Fig. 2A**). Same conventions as in **A**.

The original RML lacking the uncertainty minimization term in the VTA function sought information in proportion to reward probability (**Fig. 3Aii,** pink). The willingness to perform in the absence of instrumental incentives is explained by the first term of eq. 1, which promotes effortful actions in proportion to extrinsic instrumental or non-instrumental rewards (R). However, only the RML-C produced the enhanced sampling at 50% relative to 100% probability (**Fig. 3Aii,** black) indicating the distinct sensitivity to uncertainty shown by humans and monkeys (**Fig. 3Aiii**). The sampling patterns of the RML and RML-C were significantly different (F contrast aRML vs aRML-C: F(1,58) = 19.25, p < 0.001), with the RML-C producing higher sampling at the uncertain (50%) prior relative to the RML (post-hoc test, t(58) = 20.99, p < 0.0001) and relative to the certain (100%) prior within the RML-C (50% > 100%: t(29) = 3.96, p = 0.0004). These patterns closely reflected the NE boost, which peaked on high uncertainty trials in the RML-C but was exclusively dependent on conditioned cue value in the RML (F contrast RML vs RML-C: F(1,58) = 4.27, p < 0.05; post-hoc t-contrast RML vs RML-C for Cue1 50%: t(58) = 13.72, p < 0.0001; t-contrast within RML-C, Cue1 50% > Cue1 100%: t(29) = 4.54, p < 0.0001; **Fig. 3Aii, right**). This suggests that the intrinsic desire to reduce uncertainty depends on a distinct component in the utility function that can be signaled by NE.

To analyze whether utility based on expected surprisal could affect sampling in instrumental conditions, we implemented eq. 1 in the aRML model and tested this modified model (the aRML-C) in the informative condition in the task of instrumental information demand described in the previous section (**Fig. 2A**). The aRML-C generated similar qualitative results as the aRML but showed better performance, including a higher fraction of correct responses (**Fig. 3B**, left: aRML-C vs. aRML: F(1,58) = 6.9, p = 0.01), higher visual gain modulation (**Fig. 3B**, center; F(1,58) = 23.62, p < 0.0001), and higher rate of trial completion (**Fig. 3B**, right: aRML-C vs. aRML: F(1,58) = 8.52, p < 0.01). Thus, even when a task is governed by extrinsic rewards, performance benefits from an intrinsic drive to reduce uncertainty.

## Discussion

We propose a novel RML-based framework for understanding information demand in terms of cost-benefit tradeoffs implemented in a biologically plausible architecture. We show that this framework, previously found to capture regulation of physical effort, learning rates and working memory^7^, also naturally reproduces findings on instrumental and non-instrumental information demand without data fitting or additional parameter tuning. The results bring new insights into the computational basis of attentional enhancement in visual maps, the mechanisms of intrinsically motivated information demand, and the biological basis of algorithms that compute free energy. We discuss each in turn.

The selection of stimuli for saccades and attention has long been associated with a network of topographically organized visual and oculomotor areas that includes portions of the frontal and parietal lobes^22^. While abundant evidence shows *that* these visual areas provide selective representations that prioritize behaviorally relevant stimuli, the computational meaning of this prioritization is under debate. Our model supports the idea that prioritization emerges from interactions between the visual maps and an MPFC-centered circuit that estimates the costs and benefits of gathering information. Response enhancement in visual maps thus reflects a cognitive action – a boost initiated by the MPFC that facilitates the selection of relevant information and its linkage with actions - that entails a *cognitive cost* and is deployed selectively according to the increase in rewards it is expected to bring. This mechanism reproduces reward and uncertainty effects in LIP cells, suggesting that these effects index adaptations of visual processing optimized for obtaining a behavioral goal.

Our results shed new light on reward modulations in visual areas, particularly the longstanding controversy about their significance in area LIP^28,29^. Specifically, the results argue that, rather than representing the economic values of alternative actions as traditionally assumed^30^, the neurons reflect adjustments of cognitive states necessary to obtain a desirable outcome. One example of this dissociation is the fact that the aRML, like LIP cells, had enhanced visual responses for decisions that have prior uncertainty and require of new information relative to those that have a strong prior, even though the latter have *higher* overall value - i.e., are completed faster with higher rates of success (**Fig. 2B**, top vs bottom). A second example is the fact that the aRML, like LIP cells, generate higher responses for smaller relative to larger rewards if a boost of control is needed to keep the system engaged in the task (**Fig. 2C**). Both results clearly illustrate that, rather than encoding value *per se*, visual enhancement depends critically on the cognitive (attentional) effort entailed in obtaining a goal. This view may account for previous findings that are inconsistent with an action-value interpretation, including that LIP neurons have enhanced responses to stimuli that command a more difficult anti-saccade rather than a habitual (and more rewarding) pro-saccade^31^, stimuli that command a change in motor plan^32^, and those that indicate low-value alternatives that are avoided by a behavioral choice^33^.

We note that, consistent with available evidence^14^ and appropriate for reducing model complexity, the RML draws a sharp distinction between a role for in NE in cognitive and physical effort and the role for DA in reward-driven motivation and learning^7^. However, very little is known about the cellular basis of cognitive effort in information demand, and new evidence may well reveal the involvement of additional neuromodulators (such as modulations of visual areas by acetylcholine and/or directly by DA in addition to NE^34,35^).

A second critical question addressed by our findings concerns the intrinsic utility that animals assign to non-instrumental information, which is based on both the extrinsic rewards and epistemic value of various states. We showed that the RML accounts for these findings by postulating that the utility function has two distinct components. A reward-based component is signaled through DA responses to reward expectation^36,37^ and can energize behavior even without instrumental training^7^, accounting for the desire to reveal signals associated with positive outcomes. A second epistemic component is based on expected surprisal, which had been estimated in the original RML and incorporated in the utility function in the RML-C. With this modification, the RML reproduced empirical observations (^2^; **Fig. 3A**) and provides three specific advances. First, the result generalizes and provides a biological implementation for earlier two-factor mathematical models proposed by^2,38^. Second, it more closely captures empirical findings relative to a recent model based on the utility of anticipation^39^. According to the model of Iigaya et al., the utility of anticipation depends on reward prediction errors (RPE), such that information that is expected to produce positive RPEs is sought because it engenders a pleasant state of anticipation (“savoring”), while information producing negative RPEs is avoided as it generates an unpleasant (“dread”) state). Unlike the RML, this model does not explain why information demand in monkeys^2^ and humans^40^ *increases* and remains high even at a 100% prior reward probability - i.e., when new information is expected to have no RPE (**Fig. 3Aiii**). A third contribution of our results is showing that a sensitivity to expected surprisal can supplement the boost provided by instrumental rewards (**Fig. 3B**). Thus, individuals who are more motivated by uncertainty in non-instrumental conditions may show enhanced efficiency of information gathering in instrumental conditions - a possible new link between instrumental and non-instrumental information demand that can be investigated in future research.

A final contribution of our results is to provide a novel computational and biological interpretation of the free-energy principle, which has been discussed in a vast theoretical literature and describes exploration decisions aimed at minimizing expected free-energy^41^. The equation we propose for the utility function of the RML-C (Equation 1) approximates the equation expressing (negative) free-energy (-*F*)^42^:

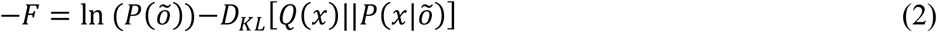

The first term in the free-energy equation indicates the log-evidence of the agent’s generative model about preferred outcomes 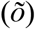 and corresponds to the extrinsic value term in our Equation 1: 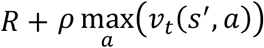. The second term in the free-energy equation is the Kullback-Leibler divergence between estimated (*Q(x)*) and true posterior 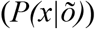 probability that the hidden cause *x* determines the outcome 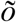, and is analogous to the second term in Equation 1, where information content is approximated by the expected unsigned prediction error: 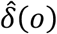. Thus, the RML-C performs approximate free-energy minimization, expressing free energy in terms of biologically plausible control theory rather than probability density functions.

This in turn leads to two significant advances. First, by implementing the free-energy principle in RL algorithms, it can spur the development of a new class of algorithms in which agents optimize free-energy, accounting for higher order types of utility that are not directly predicated on extrinsic rewards. Second, by implementing the free-energy principle in a biologically plausible architecture, the model provides a powerful tool for testing hypotheses on how motivational, executive and sensory systems are coordinated to seek information and reduce free energy.

## Methods

All the results shown in the text are based on 30 simulations of 360 trials each. These numbers are in the range of sessions and trials tested in experimental paradigms and prevent artificially inflating p-values by increasing the number of simulations. The results are not based on fitting the model to empirical data. Instead, we simulate new tasks using the same the parameters as in Silvetti et al.^7^; Table 1.

**Table 1.** RML parameters list and values for discrete model.

| Parameter | Value | Meaning | Equation |
| --- | --- | --- | --- |
| $\rho$ | 0.2 | TD-learning signal decay | 8a |
| $m$ | 0.1 | DA dynamics | 8a |
| $\tau$ | 0.6 | Softmax temperature | 4,6,10 |
| $\alpha$ | 0.3 | Learning rate hyper-parameter | 7c-d |
| $\beta$ | 0.2 | Learning rate lower bound | 7a |
| $\omega$ | 0.15 | Boosting cost | 8b |

### RML model

Here we introduce the equations governing the RML function. The software used for the simulations in this article can be downloaded from the RML GitHub repository: https://github.com/AL458/RML.

#### MPFC_Act_

The central equation in this module governs state/action value updates:

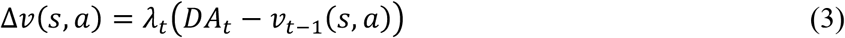

where *v(s,a)* indicates the value (outcome prediction) of a specific action *a* given a state *s*, and *DA* is the environmental outcome, here interpreted as dopamine signal afferent from VTA (Equation 8). The learning rate parameter *λ*, governing the flexibility of value update, is computed by Equation 7a.

The probability of selecting action *a*, conditioned to state *s*, is given by a softmax function σ, whose arguments are the state/action values *v* discounted by state/action costs *C* and temperature *τ*

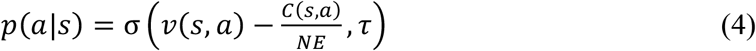

Matrix *C* assigns a cost to each state/action couple, for example energy depletion consequent to climbing an obstacle. *C* is discounted by norepinephrine afferents from LC (*NE*), which is itself controlled by the MPFC_Boost_ module, via parameter *b* (Equation 6).

#### MPFC_Boost_

This module controls the parameters for cost and reward signals in equations 3-4 (MPFC_Act_), via modulation of VTA and LC activity (boosting catecholamines). This is implemented by selecting the modulatory signal *b* (*boost signal*), by RL-based decision-making. The MPFC_Boost_ updates the boost values *v_B_*(*s, b*), via the equation:

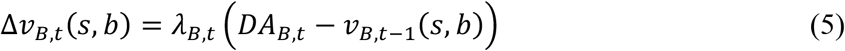

Equation 5 represents the value update of boosting level *b* in the environmental state *s*. The MPFC_Boost_ submodule selects boosting actions probabilistically, based on expected values *v_B_* and temperature *τ*

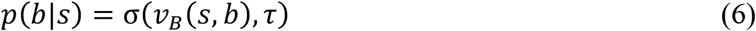

Here, we imposed ***NE = b***, for all the equations involving the norepinephrine output for effort implementation.

#### Control over learning rate

The learning rate parameters in the two MPFC modules (*λ* and *λ_B_*) are optimized online (i.e. while the model interacts with the environment) as a function of both uncertainty and volatility. Optimization of *λ* and *λ_B_* solves the trade-off between stability and plasticity, increasing learning when the environment changes and lowering it when the environment is simply noisy. *λ* and *λ_B_* are computed as the ratio between the estimated variance of state/action-value 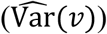 over the estimated squared prediction error 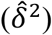 (Kalman gain approximation^43^; for simplicity we here indicate only *λ*)

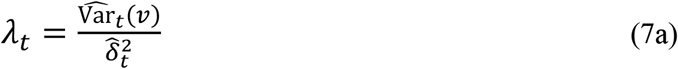

With *β* ≤ *λ* ≤ 1 (*β* is a free parameter indicating the minimal *λ* value), to ensure numerical stability. The process variance is given by:

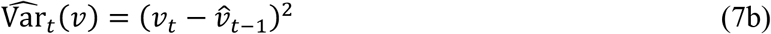

where 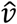 is the estimate of *v*, obtained by low-pass filtering tuned by the hyperparameter *α*:

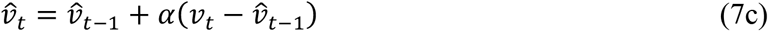

The same low-pass filter is applied to the prediction error signal (*δ*) to obtain a running estimation of total variance 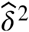, which corresponds to the squared estimate of unsigned prediction error:

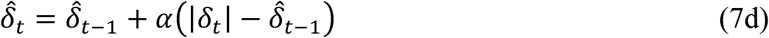

Equations 7a-d are implemented independently for each of the two MPFC modules, so that each module has its own learning rate parameter (*λ* or *λ_B_*).

**VTA** The VTA module provides outcome-related signal *DA* to both MPFC modules, either for action selection directed toward the environment (by MPFC_Act_) or for boosting-level selection (by MPFC_Boost_) directed to the brainstem catecholamine nuclei. Equation 8a below defines the DA signal to the MPFC_Act_:

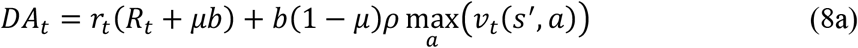

Where, *r* is a binary variable indicating the presence of reward signal, and *R* is a real number variable indicating reward magnitude. Parameter *ρ* is the TD discount factor, while parameter *μ* is a scaling factor distributing the modulation *b* between primary (first term of the equation) and non-primary (second term) reward.

The VTA signal afferent to the MPFC_Boost_ is described by the following equation:

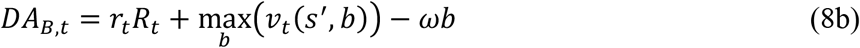

where *ω* is a parameter defining the cost of catecholamine boosting.

### The aRML hybrid model

We used a recurrent basis function network with continuous attractor dynamics^18^ to simulate LIP neurons (**Figure 1b**). The LIP network consisted of three noisy input layers that were also the output ones (recurrent dynamics): a layer coding visual information in the eye-centered frame of reference, a layer coding for the horizontal position of the eye in the orbit and a layer coding for the visual information in horizontal head-centered frame of reference. All the layers were connected, with symmetric weights, to a hidden 2-dimensional layer (basis function layer) representing a 2D map that integrated information from different frames of reference and encoding the direction of the future saccadic movement (shown in Figure 1c). At the beginning of each trial, gaze was on fixation point and the cue appeared either in the left or in the right hemifield. The 2D map of LIP neurons encoded the direction of the future saccade toward the cue. A full simulation of visual processing would have required representing both LIP (for saccade planning) and MT (for motion discrimination). Because such a complexity level was unnecessary to test our model, here we linked directly the cue decoding to the precision of saccadic movements (Equation 14), without explicitly simulating the MT area for motion decoding. In order to do so, we assigned the meaning of motion direction to the eye-centered dimension, i.e. the eye-centered visual information “up” or “down” indicates movement direction (respectively “up” or “down”). Equations and parameters of the LIP network are reported in the supplementary material of the original article where the network was described^18^, here we did not introduce any modification.

#### Interface RML-LIP

The RML modulated the output of the LIP hidden layer by means of *NE* signal. Given that the LIP hidden layer is a 2D map (*A*), each neural units is indexed by a couple (*l,m*), where *l* indexes the influence of the eye position, while *m* of the visual information. If we define the activity of the unit (*l,m*) as *A_lm_*, the RML modulated this activity by the following equation:

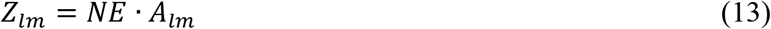

Where *NE* is the norepinephrine output from the RML (scalar). Then, the state-action value updated by the information provided by *Z* is:

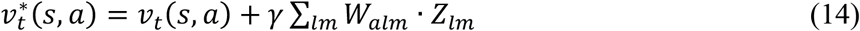

Where *v*\* is the biased state-action value and *W* is a weight matrix for the linear transformation from *Z* map to state-action value map (*a* ∈ [1,2]) and *γ* = 10^-3^ is a scaling parameter; *v*\* is then passed as an argument to Equation 4 for action selection. The weights matrix *W* is defined as follows:

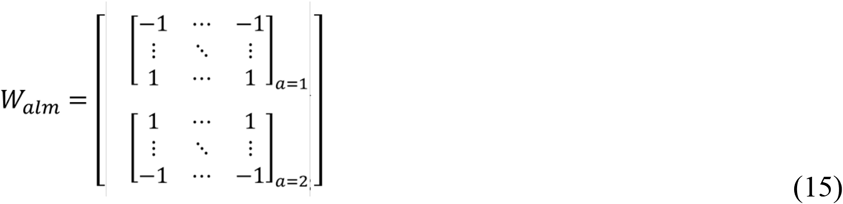

This guarantees opponent coding of cue direction by the upper and the lower half of the basis function layer *Z_lm_*.

To study the effect of task difficulty in the Instrumental Information Sampling Task (Figure 3C), we increased difficulty by decreasing the *γ* parameter (10^-3^ ≤ *γ* ≤ 6 · 10^-4^) to make progressively weaker the choice bias evoked by the cue (Equation 14). Fisher information 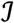 provided to the RML by the bias term in Equation 14 was computed as a function of *γ*:

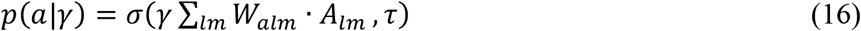

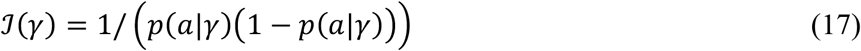

Where *σ* is the softmax function providing the probability *p(a)* that the cue alone (without NE modulation) could bias the action toward the correct action *a*.

### Equations relative to Non-Instrumental Sampling Task

Analogously to expected free-energy in Active Inference algorithms^42^, decision-making was aimed at maximizing *DA* input expected alongside the path integral of the decision tree relative to each possible policy. The value *V* relative to policy *π*was computed by the following path integral:

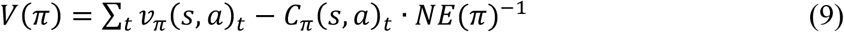

Where *v_π_(s, a)_t_* and *C_π_(s, a)_t_* represent respectively the sequence of state-action values (Equation 3) and state-action costs encountered by following the policy *π. NE(π)* is the norepinephrine level selected for the policy *π*

The best policy was then selected probabilistically, by means of a softmax function σ, with temperature τ:

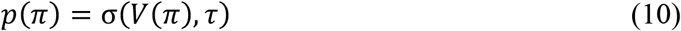

*DA* signal (substituting Equation 8a) was computed as:

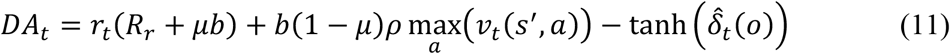

Where 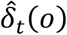 is the expected surprisal expressed as the expected unsigned prediction error for outcome *o*, computed by Equation 7d. The hyperbolic tangent function guarantees the information surprise term belongs to the interval [0,1]. Equation 11 has been represented in a simplified version (Equation 1) in the Results section.

Finally, *DA*_B_ (substituting Equation 8b) included the information surprise component, becoming expressed by the following equation:

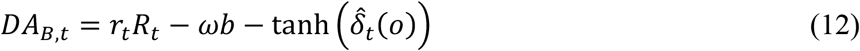

Where 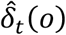 is defined for Equation 11.

## Acknowledgments

The research described in this paper was supported by a National Eye Institute Grant R01EY025158 to JG, a National Institute of Mental Health R01 MH-098039 to JG, scholarships from Knud Højgaards Fond, Reinholdt W. Jorck og Hustrus Fond, and Viet-Jacobsens Fond to MH

